# Scaling diversity, abundance and energy: The Equilibrium Theory of Biodiversity Dynamics

**DOI:** 10.1101/2024.12.04.626563

**Authors:** David Storch, Jordan G. Okie

## Abstract

This preprint of a book chapter presents the newly proposed Equilibrium Theory of Biodiversity Dynamics (ETBD), whose aim is to conceptualize large-scale dynamics of species richness via addressing the population size-dependence of speciation and extinction rates, the resulting diversity-dependence of these rates, and their modulation by the environment. It provides the most general framework for understanding large-scale biodiversity patterns such as the latitudinal diversity gradient (LDG) and temporal patterns of biodiversity changes. The theory has been published elsewhere in its full form that includes all the derivations (Okie & Storch 2004, American Naturalist https://doi.org/10.1086/733103), but here it is presented in a simpler and user-friendly way, focusing on its major implications comprising macroecological scaling relationships between energy (or resource) availability, species richness and community abundance.

## Introduction

A central goal of ecology is to understand patterns of biological diversity and abundance. The field of macroecology, which sprung up in the 90s, has fostered extensive descriptions and analyses of many of these patterns, including patterns in diversity across spatial and temporal scales, taxa, and environmental gradients, and aimed to understand, among other things, the links between species richness, abundances, body sizes and flows of energy through ecosystems (Gaston and Blackburn, 2000). Today, however, we still lack a consensus about the origin of most of these patterns, including perhaps the most famous and thoroughly studied one, the latitudinal diversity gradient (LDG) (Pontarp et al., 2019). There are many factors proposed to play a role in the origin of large-scale diversity patterns, and a few theories that pretend to be general, but ecologists have not been able to reconcile these differing approaches. A similar lack of consensus comprises patterns of species and community abundance along environmetal gradients, as well as the nature of changes in the diversity, abundance and biomass of communities through time.

Here we argue that the progress in understanding macroecological biodiversity and abundance patterns has been impeded by a lack of appropriate theory addressing the coupled dynamics of abundance and diversity. Recently, we have been working on a theory to address this shortcoming of the current body of macroecological theory, which we call the Equilibrium Theory of Biodiversity Dynamics (ETBD)(Okie and Storch, 2024; Storch et al., 2018, 2022; Storch and Okie, 2019). We first describe the motivation of the theory based on what we have learned about macroecological biodiversity patterns in the last few decades. Then we describe the core of the theory and show how it can be used to predict the variation of diversity along environmental gradients and scaling relationships between energy availability, species richness and community abundance. We argue that ETBD represents a proper general framework for understanding biodiversity dynamics and large-scale temporal and spatial diversity patterns, as it addresses the coupled dynamics of species richness and community abundance modulated by energy or resource availability and species origination and extinction dynamics.

### What have we learned about spatial and temporal species diversity patterns

Regardless of the mentioned lack of consensus concerning the origin of major large-scale biodiversity patterns, the three last decades of research brought several insights that can be considered as relatively settled:

1. Terrestrial diversity is tighly related to climatic variables, namely temperature and precipitation (Field et al., 2009), ecosystem net primary productivity (NPP) being typically the best single correlate of species richness (*S*) of consumers at large geographic scales. The amount of resources thus seems to be a major determinant of diversity patterns (Bohdalková et al., 2021), at least for terrestrial ecosystems (Fig. 1A).
2. Patterns of community abundance (*J*) are generally weaker than species richness patterns, and even though community abundance often positively correlates with species richness, it typically does not follow environmental gradients in the same way as species richness (Currie et al., 2004; Storch et al., 2018). Consequently, mean per species population abundance 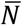, which is equal to J/S, typically varies along environmental gradients (Fig. 1B).
3. Species richness patterns for higher taxonomic levels have a tendency to converge regadless of historical circumstances (Hawkins et al., 2012), but species richness is often not correlated or is negatively correlated with the measures of speciation rate (Igea and Tanentzap, 2020; Machac, 2020; Rabosky et al., 2018).
4. Over both ecological and geological time scales, diversity varies with environmental conditions, but when conditions remain comparatively constant, diversity is rather stable (Brown et al., 2001; Close et al., 2020, 2019). Such stability seems to be maintined by negative diversity-dependence – across geological timescales, higher diversity has been shown to lead to decreasing origination and/or increasing extinction rates (Rineau et al., 2022).

**Fig. 1.**
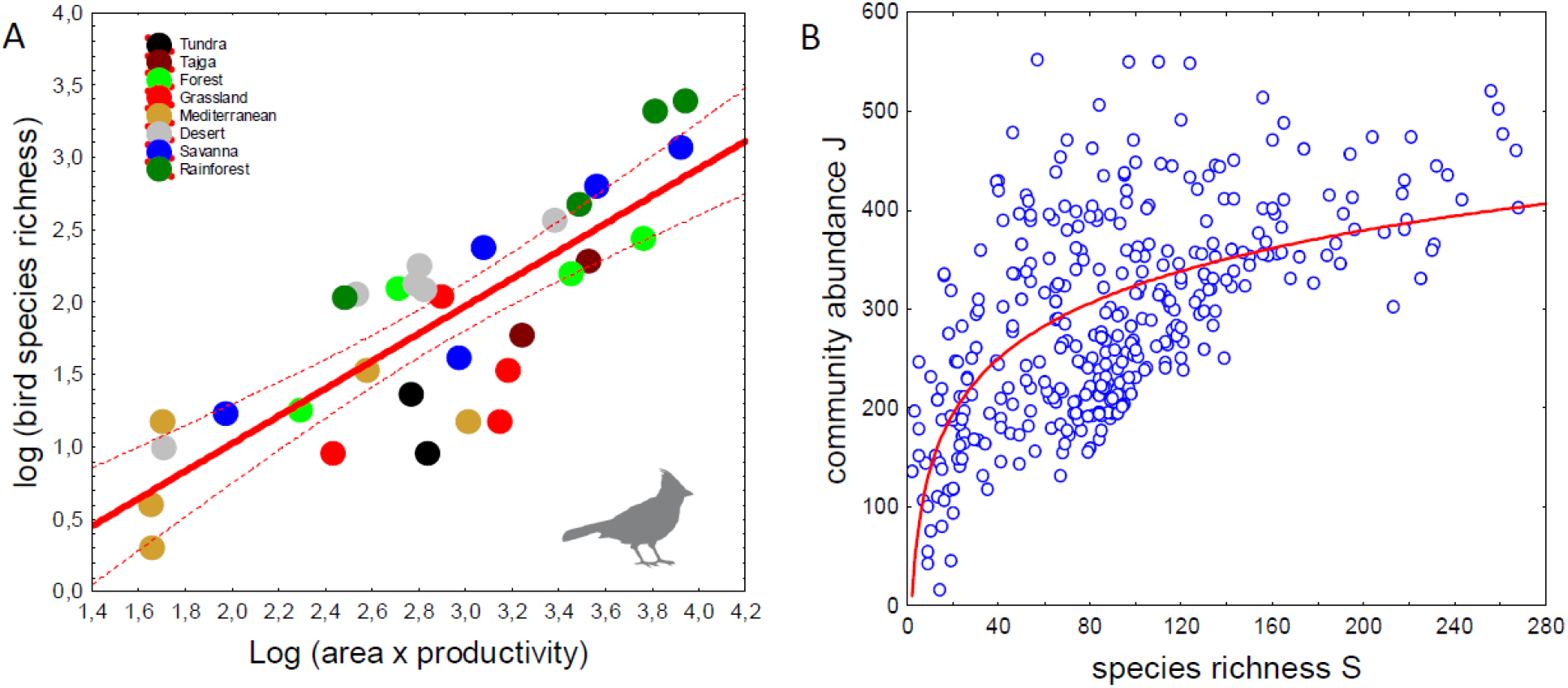
Illustrative examples of scaling relationships between energy availability, species richness and community abundance. (A) Empirical scaling relationship between the total mass of organic carbon produced per unit time by the plants of a biome (i.e. total net primary production, NPP, which is directly related to energy availability for consumers and is caclulated as NPP x biome area) and bird species richness across different biomes within individual biogeographic realms (data from Jetz and Fine, 2012). It is approximately linear in the log-log scale, so it can be reasonably well approximated by a power-law. (B) The relationship between species richness *S* and community abundance *J* for 0.1 ha forest plots (Gentry plots; data from Šímová et al., 2011). The axes are not log-transformed to show the non-linearity of this relationship (fitted by logarithmic function in this particular case).

None of these findings can unequivocally provide a clue to the major causes of diversity patterns like the LDG. However, in the light of these findings, some classical explanations of large-scale diversity variation seem obsolete. For instance, many hypotheses of the LDG propose that higher diversity in the tropics is due to higher speciation rates (Pontarp et al., 2019); however, these hypotheses are refuted by the frequent observation of a decoupling between diversity and speciation patterns (point 3; Rabosky et al., 2018). In contrast, historical explanations based on the idea that historically older, larger and more stable regions had more time to accumulate species, may be valid for some relatively smaller clades, but they are not entirely compatible with the finding that diversity dynamics is regulated by negative diversity-dependence (point 4) nor with the convergence of diversity patterns across taxa regardless of different histories (point 3).

Taken together, it seems most probable that major diversity patterns reflect, at least to some extent, equilibria of diversity dynamics or region-specific diversity limits (Storch & Okie 2019). However, these limits usually do not appear to be directly driven by a limited number of individuals (i.e. the more-individuals hypothesis, Gaston 2000), as *J* varies more weakly than species richness along large-scale environmental gradients (Storch et al., 2018; see point 2). It is thus useful to develop the idea of these limits or diversity equilibria more formally. This is the major purpose of ETBD.

### Cornerstones of ETBD

ETBD is based on two intuitive assumptions that may be considered unequivocal principles of ecology (Storch et al. 2022, Okie & Storch, 2024). First, diversity dynamics is modulated by negative diversity-dependence, which universally follows from the population size-dependencies of extinction and origination rates. The basis of this principle is straightforward: for a given total amount of resources, increasing species richness must lead to progressively decreasing population sizes. Eventually, if species richness is very high, many populations are so small that their extinction rates are higher than their origination rates, and consequently diversity has a tendency to decrease. Conversely, when species richness is low, population sizes are large, on average, and consequently more species have lower probabilities of extinction than origination, leading to species richness increasing until the total rate of extinction is balanced by the total rate of origination. Note that the equilibrium diversity thus depends not only on the total amount of resources but also on the rates themselves and how they depend on population size (Storch & Okie, 2019).

The second assumption of ETBD is that total community abundance is not directly driven by the properties of the environment, but arises from the combination of these properties - namely the total amount of available resources, such as energy - and species richness itself. The idea stems from the observation that more species can utilize more resources, and although there is a ceiling for this increase given by total energy availability, the addition of species to a community often leads to the utilization of previously unutilized resources and the more efficient conversion of resources into standing community biomass (Cardinale et al., 2006; Loreau, 1998; Nijs and Roy, 2000; O’Connor et al., 2017). We call this the biodiversity effect on community abundance (BECA), which is one of the specific manifestations of the biodiversity-ecosystem function relationship (BEF) that has been relatively well-documented (Cardinale et al., 2006; O’Connor et al., 2017; Tilman et al., 2001). ETBD thus assumes population sizes are ultimately limited by competition for shared resources, but allows for at least partial niche complementarity and an expansion of total niche space with an increase in the number of species.

These two essential assumptions lead to an interesting dynamical interlinkage between species richness and community abundance, as equilibrium species richness is regulated by community abundance, but species richness in turn itself affects community abundance due to the biodiversity effect on community abundance (BECA). To address this interlinkage, it is necessary to conceptualize the dynamics as a movement in the two-dimensional state (or phase) space of *S* and *J* (Fig. 2). If we plot community abundance *J* against species richness *S*, there must be at least one diagonal line that depicts all potential species richness equilibria *Ŝ* in which origination and extinction rates are balanced. The reason is that, everything else being equal, there is a mean equilibrium population size 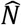 at which the rates are balanced, and the equilibrium diagonal line consequently follows equation 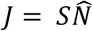. We call this diagonal line the S-nullcline. Communities located right of their S-nullcline have lower average abundances than 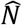, so that their extinction rates are higher than their origination rates, leading to movements in the state space towards their S-nullcline. The opposite holds for communities located to the left of their S-nullcline; the S-nullcline thus represent stable diversity equilibria.

**Fig. 2.**
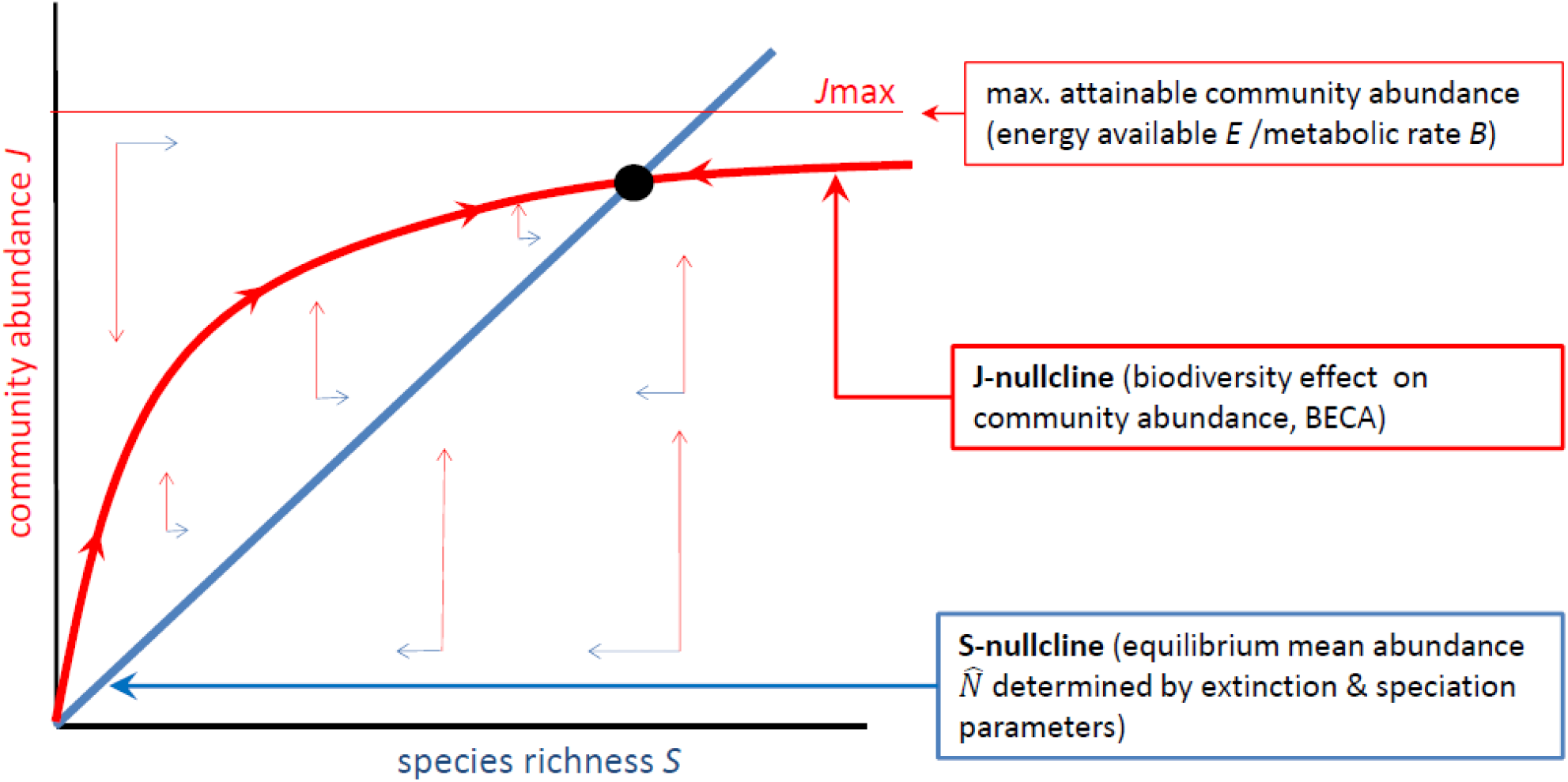
The core of the Equilibrium Theory of Biodiversity Dynamics (ETBD). According to the theory, the dynamics of species richness (*S*) and total community abundance (*J*) is coupled and can be modeled as a movement in a two-dimensional S-J state space. The blue line depicts the S-nullcline, which is characterized by equal total speciation and extinction rates; all the points along the S-nullcline thus represent equilibria of species richness. The reason is that for any community, there is an equilibrium mean abundance 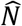 where speciation and extinction rates are balanced – smaller populations have a higher probability of extinction than speciation, while larger populations have a higher probability of speciation than extinction. Communities located to the right of the S-nullcline have mean abundance J/S lower than 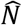. Consequently, their extinction rates prevail over speciation rates and they have a tendency to move left (decrease their species richness), while communities to the left of the S-nullcline move right (blue vectors). The S-nullcline thus represents stable equilibria of species richness - it is an attractor nullcline. ETBD additionally addresses the possibility that species richness can affect equilibrium community abundance *Ĵ*, e.g., because more species are generally able to utilize more resources (there is a biodiversity effect on community abundance, BECA). The J-nullcline that characterizes this effect (red line) can have various forms, but here we assume that it is an increasing function with a ceiling, given by limited total energy availability *E* and mean individual metabolic rate *B*, so that *J*_*max*_ = *E*/*B*. Since population dynamics leading to the utilization of all resources available for given set of species is considerably faster than evolutionary dynamics of speciation and extinction, the movement in the vertical (abundance) direction (red vectors) is much faster than in the horizontal (species richness) direction. Consequently, a community has a tendency to reach relatively quickly the J-nullcline, and then move more slowly along it towards the equilibrium point (black dot), given by the intersection of the S-nullcline and J-nullcline. This equilibrium point is thus determined by (1) the balance of speciation and extinction that determines the slope of the S-nullcline, (2) E/B, which determines the ceiling of the J-nullcline, and (3) the overall shape of the J-nullcline.

The S-nullcline shows the values of equilibrium species richness present for different values of *J*, so to determine the attractor of the dynamics (the stable equilibrium point), we need to also specify equilibrium *J* in the phase space; in other words, the J-nullcline. If species richness has no effect on equilibrium *J*, then the J-nullcline is a horizontal line, as implicitly assumed in other theories of biodiversity dynamics (e.g., Hubbell’s neutral theory; Hubbell, 2001). However, if species richness has a positive effect on community abundance, as is generally found across taxa and ecosystems (Cardinale et al., 2006; O’Connor et al., 2017), then the J-nullcline is a positive, increasing curve. It can have various forms, but here we assume for simplicity that *Ĵ* increases with *S* (for the reasons mentioned above), but with a decelerating rate due to the limited total amount of resources (for more complex shapes of J-nullcline see Okie & Storch, 2024). This J-nullcline represents an attractor nullcline, as *J* above the nullcline is not sustainable in the long term (species cannot utilize more resources than what is allowed by *S* and the total amount of resources) and all *J* below the J-nullcline have a tendency to move towards the nullcline due to population dynamics approaching the carrying capacity. The dynamics within the S-J state space thus eventually leads to a stable equilibrium point located at the intersection of the S-nullcline and J-nullcline (Fig. 2).

### Scaling equilibrium species richness and community abundance along environmental gradients

Hereafter we consider large-scale gradients of environmental variables that can affect respective processes within large regions or provinces in which speciation rate is much more important than colonization rate (Rosenzweig, 1995, Storch et al., 2018). We will thus neglect dispersal and colonization dynamics, and instead of origination rate we will speak specifically about speciation rate. It can be seen from Fig. 2 that the position of diversity equilibria within the S-J state space are determined by three factors: (1) the overall levels and population size-dependencies of speciation and extinction rates that determine 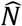, and thus the slope of the S-nullcline, (2) the total energy or resource availability that determines the ceiling of J-nullcline and thus its overall level, and (3) the shape of the J-nullcline. All these factors can vary across environmental gradients, affecting the scaling of equilibrium *S* and *J* along the gradients (Fig. 3).

**Fig. 3.**
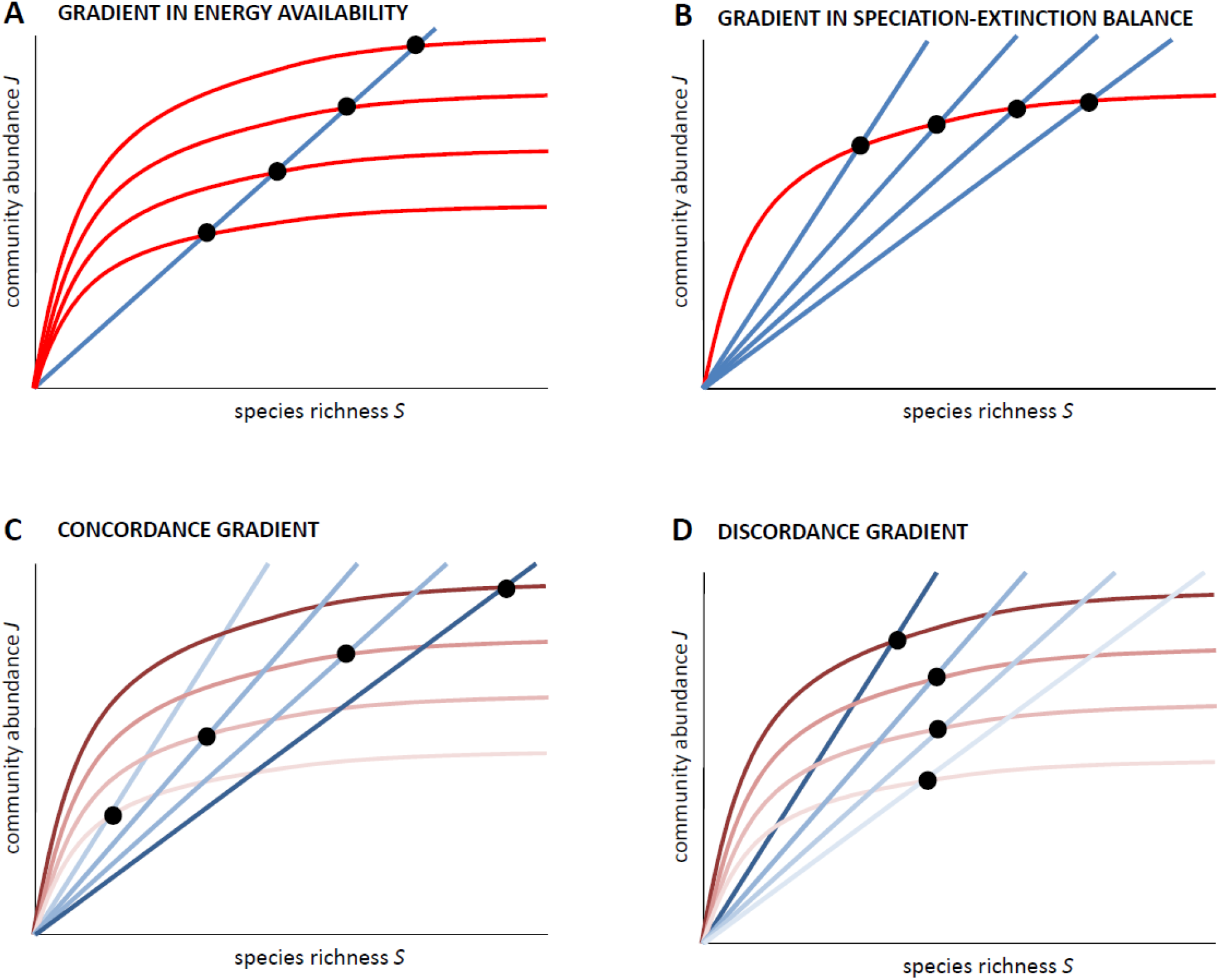
Illustration of how scaling relationships between equilibrium *S* and *J* emerge in the four fundametal categories of diversity gradients. (A) If only energy availability *E* or mean metabolic rate *B* vary along a gradient, leading to pure variation of *J*_*max*_, all equilibria (black dots) lie on one invariant S-nullcline (blue line), and species richness and community abundance are thus predicted to be proportional to each other. (B) In contrast, if the J-nullcline (red line) does not change along a gradient and only factors driving the speciation-extinction balance vary, leading to varying S-nullclines, all equilibria follow the J-nullcline; the S-J scaling can be thus non-linear, depending on the shape of the J-nullcline. (C) If both these factors vary along a gradient, but the factors increasing speciation or decreasing extinction are positively correlated to E/B (here depicted by increasingly dark colours of S-nullclines and J-nullclines along a gradient), we speak about *concordance gradients*, as the environmental gradient fosters increasing equilibrium species richness due to both of the effects. *S* thus scales positively and non-linearly with *J*, and also with E/B. (C) in contrast, if the factors that increase speciation or decrease extinction rates are negatively correlated with E/B, we speak about *discordance gradients*, and S-J scaling, as well as the scaling of species richness with energy availability, may have various forms, including a hump-shaped relationship between S and J and between energy availability and species richness. Assuming particular functions for all the relationships and the level of concordance/discordance of the gradient, we can quantitatively predict all the scaling relationships (Box 1, Figure 4).

Here we will not consider the third effect, i.e. changes in the J-nullcline shape, as there are no clear expectations about systematic changes of the J-nullcline shape along any environmental gradient. In contrast, there are reasons to expect systematic changes of the two first factors that lead to particular predictions concerning the scaling of equilibrium *J* and *S* along environmental gradients and to each other. Fig. 3A shows that if only resource/energy availability changes along a gradient, a linear scaling between equilibrium *J* and *S* is expected, as all the communities lie on an unchanging S-nullcline. In contrast, if only the speciation-extinction balance changes along the gradient, i.e. the slope of the S-nullcline changes, then the scaling between equilibrium *J* and *S* follows the J-nullcline instead and so is expected to be non-linear (Fig. 3B).

The situation becomes more complex when both resource/energy availability and speciation-extinction balance vary simultaneously along a gradient, such as with latitude. In this situation, the scaling relationships depend on the covariance of these two factors along the gradient, i.e. whether they are concordant (Fig. 3C) or discordant (Fig. 3D), leading to a *concordance gradient* or *discordance gradient*, respectively. Discordance gradients may be typical, for instance, for the oceans, in which productivity is frequently higher at higher latitudes, while extinction rates may be lower in the tropics. In such cases, *J* may even decrease with increasing *S* or the relationship can be hump-shaped or U-shaped. The reason is that when resource availability is positively associated with extinction levels or negatively associated with speciation levels, lower diversity equilibria can emerge regardless of an augmentation in *J* resulting from the higher resource availability.

### Quantitative predictions of ETBD

Over a given range of *S*, the J-nullcline can be mathematized as a power-law function of *S* (e.g., Liang et al., 2016; Okie and Storch, 2024) such that

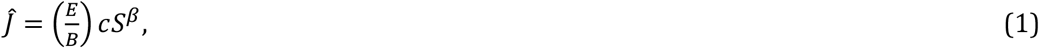

Where *Ĵ* is the equilibrium community abundance (the *J* at which dJ/dt = 0), *c* sets the fraction of the maximum available energy *E* used by a one-species community, and *β* quantifies the biodiversity effect on community abundance (BECA). A thermodynamic constraint to Equation 1 is that *Ĵ* cannot be greater than maximum community abundance *J*_*max*_, which is determined by the maximum available energy *E* divided by average individual metabolic rate *B* (i.e., *E*/*B*).

A power law model for the BECA relationship has empirical support from studies of the effect of diversity on community biomass production, standing biomass, and resource consumption (Liang et al., 2016; O’Connor et al., 2017; Reich et al., 2012) and some theoretical basis (Liang et al., 2016, 2015; Mora et al., 2014; Okie and Storch, 2024). When the *β* is between 0 and 1, the J-nullcline is a decelerating cuve, implying that competition (niche overlap) along with some level of niche complementarity mediate the biodiversity effect on community abundance. In general, the closer *β* is to zero, the more intense the competition for resources. Given the importance of competition in shaping communities and that empirically *β* averages around 0.26 across a variety of taxa and ecosystems (O’Connor et al., 2017), here we focus on predictions for 0 ≤ *β*< 1 (see Okie and Storch, 2024, for predictions concerning *β*>1).

ETBD focuses on two fundamental gradients along which biodiversity equilibria may change: gradients in energy availability and gradients in the extinction-speciation balance. Each gradient exhibits a different scaling of *S* with *J*. Along gradients in which the slope of the S-nullcline (i.e. speciation-extinction balance) is changing, ETBD predicts (see also Fig 3B)

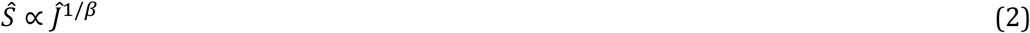

In contrast, along gradients in maximum energy availability *E*, maximum community abundance (E/B), and individual metabolic rate *B*, *Ŝ* is expected to scale linearly with *Ĵ* (Fig. 3A):

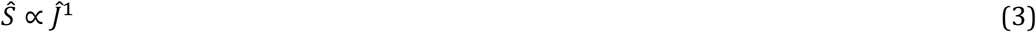

Along energy gradients, ETBD predicts that

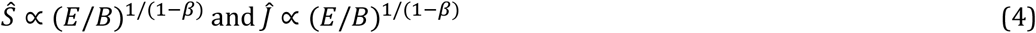

In log-log plots of *Ŝ*, *Ĵ*, and *E*, straight lines are expected with slopes equal to the exponents in the equations. These equations show that the scaling exponent (*β*) quantifying the biodiversity effect on community abundance (BECA) shapes the scaling of *Ŝ*, *Ĵ*, and *E*/*B*.

Remarkably, when competition with niche complementarity govern the J-nullcline (0 < *β* < 1), as is generally expected, *Ŝ* and *Ĵ* increase superlinearly (disproportionately) with *E*/*B*, and *Ŝ* scales superlinearly with *Ĵ* along speciation/extinction gradients. *β* tends to have relatively small values, averaging 0.26 across taxa and ecosystems (O’Connor et al., 2017), so *Ŝ* and *Ĵ* are generally expected to increase moderately superlinearly with E/B as approximately across taxa and ecosystems and *Ŝ* is expected to scale highly superlinearly with *Ĵ* along speciation-extinction gradients as approximately *Ŝ* ∝ *Ĵ*^3.85^ These predictions could be made more precise for specific taxa or ecosystems as our understanding of the J-nullcline advances.

ETBD also makes predictions for the scaling of *Ŝ* with *Ĵ* along complex gradients in which the extinction-speciation balance (i.e. equilibrium mean abundance 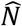) and energy availability (E/B) are varying simultaneously along the gradient (Box 1). In a concordance gradient, 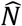 decreases with increasing energy availabilty, leading to the scaling of S with J having an exponent between 1 and 1/*β*—that is, between the scaling expected for a pure resource-availability gradient and the scaling expected for a purely speciation-extinction gradient (see Fig. 3B). In contrast, in a discordance gradient, 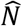 increases with increasing energy availabilty, such that at higher resource availability, speciation is lower or extinction is higher (see Fig. 3C). In such a situation, a great variety of positive and negative S-J relationships can occur, depending on *β* and whether the speciation/extinction balance versus *E*/*B* is varying most along the gradient. In a perfectly discordant gradient, the drivers of diversity vary completely opposingly leading to no change in species richness along the gradient, but there is still a change in *J* along the gradient that is determined by *β*.

ETBD also makes predictions for how *Ŝ* and *Ĵ* should change as a function of an environmental variable G along such comlex gradients. Okie and Storch (2024) made predictions for the two most general scenarios, which apply to a variety of situations: gradients in which *E*/*B* and the speciation/extinction balance (i.e., equilibrium mean abundance 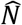) are associated with some environmental variable as (1) a power function or (2) an exponential function (Box 1). Using this framework, predictions can also be readily derived for particular gradients in which changes in *E*/*B* and the speciation/extinction balance are not well characterized by an exponential or power power law function, for example, due to E/B saturating along the gradient. Taken together, ETBD shows how species richness and abundance patterns along environmental/geographic gradients depend on (1) the degree of concordance or discordance between two major drivers of diversity variation, namely speciation/extinction rates and resource availability, (2) the scaling of the biodiversity effect on community abundance (BECA), (3) and how equilibrium mean abundance and energy availability are changing along the environmental gradients.

## Discussion

The Equilibrium Theory of Biodiversity Dynamics is able to predict a wide range of patterns of species richness and community abundance along environmental gradients under particular settings, based on general and intuitive assumptions. In contrast to other theories addressing biodiversity dynamics like the neutral theory (Hubbell 2021), it does not assume constant and species richness-independent community abundance, and allows for a wide spectrum of speciation and extinction dynamics and their population size-dependencies. It thus represents a robust foundation for understanding large-scale diversity patterns. It reconciles different views on diversity dynamics, namely the idea of diversity limits and the notion that species can be added to a community so that the overall niche space increases.

The scaling predictions of ETBD remain to be tested, as ETBD predicts different scaling relationships under different settings, which are often difficult to evaluate. However, evaluating patterns of diversity simultaneously with patterns of community abundance along environmental gradients can illuminate the role of various factors and processes responsible for spatial diversity gradients. Specifically, Okie & Storch (2024) have shown that global patterns in tree diversity and abundance were remarkably well predicted by assuming they are purely related to variation in the speciation-extinction balance rather than energy availability across latitudes. Also, scaling relationships predicted by the ETBD may be useful in predicting biodiversity changes in the Anthropocene. For instance, the predicted superlinear scaling of *Ŝ* and *Ĵ* with total resource availability indicates that a decrease in resource availability, e.g., as resulting from habitat loss and human harvesting of biomass, could lead to a disproportionate loss of species richness (Storch et al., 2022).

That said, there are several important caveats and nuances. First, and most importantly, ETBD is a theory concerning large-scale patterns and dynamics, in which the basic spatial units (communities) are large provinces like ecoregions or biomes within geographic realms that are relatively independent of each other. The reason is that ETBD does not address dispersal and colonization dynamics, which are of increasing importance to diversity patterns at smaller spatial scales. These processes have population size-dependencies and diversity-dependencies that are more difficult to evaluate and more idiosyncratic. Dispersal may homogenize communities, distorting the scaling relationships predicted by the theory. Communities that represent smaller samples of the ecoregions are expected to reveal patterns of abundance along environmental gradients which are similar to the patterns predicted for large communities by ETBD, since abundance scales linearly with area (smaller areas have proportionally lower community abundance). However, species richness patterns may differ for smaller areas due to nonlinearities in the species-area relationship (Storch, 2016). Addressing these spatial scale-dependencies and the role of colonization and dispersal is a challenge for future research.

Second, equilibrium mean abundance 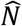 could also vary along a gradient if the form of the species abundance distribution (SAD) varies along a gradient. The form of the SAD could change, for example, due to changes in the assembly dynamics along the gradient, such as the relative balance of niche-based and neutral processes (Pueyo et al., 2007). Although we are not aware of any strong a priori reasons to expect systematic changes in the form of the SAD along typically studied large-scale gradients, if such changes were observed, they can be incorporated into the theory and used for testing its predictions.

Last but not least, in reality, many communities may be out of equilibrium, and disequilibrial communities could reveal different scaling patterns than those predicted by the theory. There is evidence that many communities are away from their diversity equilibria, especially during periods of substantial environmental change (Šímová et al. 2024). Such rapid change characterizes the current period of the Anthropocene, and the scaling relationships may be thus subject to considerable variation. Still, the equilibrium situations represent attractors of the dynamics and thus provide baseline predictions of observed patterns of diversity and abundance related to environmental variation.

## Acknowledgements

The study was supported by the Czech Science Foundation (grant number 20-29554X).

## Box 1

**Scaling predictions for complex environmental gradients**

ETBD can be used to make predictions for complex gradients in which the extinction-speciation balance (i.e. equilibrium mean abundance 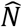) and energy availability (E/B) are varying simultaneously along an environmental gradient *G*. Okie and Storch (2024) consider the two most general scenarios: (1) *E*/*B* and the speciation/extinction balance (i.e., equilibrium mean abundance 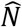) are associated with some environmental variable *G* as power functions, such that 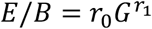 and 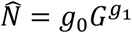, where *r*_0_ and *g*_0_ are normalization coefficients and *r*_1_ and *g*_1_ are scaling exponents quantifying the effects of *G*; or (2) *E*/*B* and the speciation/extinction balance (i.e., equilibrium mean abundance 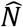) are associated with some environmental variable *G* as an exponential function, such that 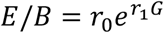 and 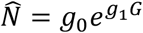. When studying a particular gradient, the choice of the power or exponential functions should be made based on theoretical or practical reasons. For instance, researchers typically consider species richness to vary exponentially with temperature and latitude (Currie et al., 2004; Storch, 2012), both because temperature has an exponential effect on biological rates (Brown et al., 2004) and because scatterplots of log richness versus temperature and latitude tend to be more statistically well-behaved compared to log-log or untransformed plots of these variables. Regardless, in both gradient scenarios, ETBD predicts:

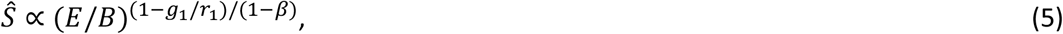

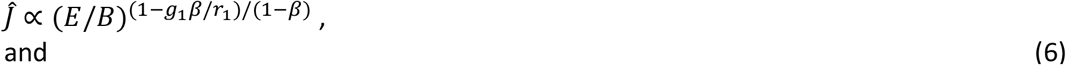

and

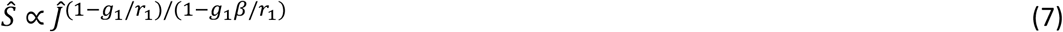

Along a purely resource-driven gradient, equilibrium mean abundance doesn’t vary and so and *g*_*1*_ = 0 and *Ŝ* ∝*Ĵ*^1^, as expected by graphical analysis of the nullclines in Figure 3. Along a purely speciation/extinction-driven gradient, resource availability doesn’t vary and so *r*_*1*_ = 0 and *Ŝ* ∝ *Ĵ* ^1/*β*^. When 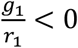, the two drivers (the positive effects of the speciation/extinction balance and resource availability on diversity) are concordant along the gradient, leading to S-J scaling being between *Ŝ*∝*Ĵ*^1^ and *Ŝ*∝*Ĵ* ^1/*β*^ When 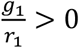, the effects are discordant, leading to a variety of potential negative and positive scalings, depending on the relative values of *g*_*1*_ and *r*_*1*_ (Fig. 4) In a perfectly discordant gradient, 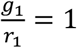resulting in no change in diversity along the gradient.

**Fig. 4.**
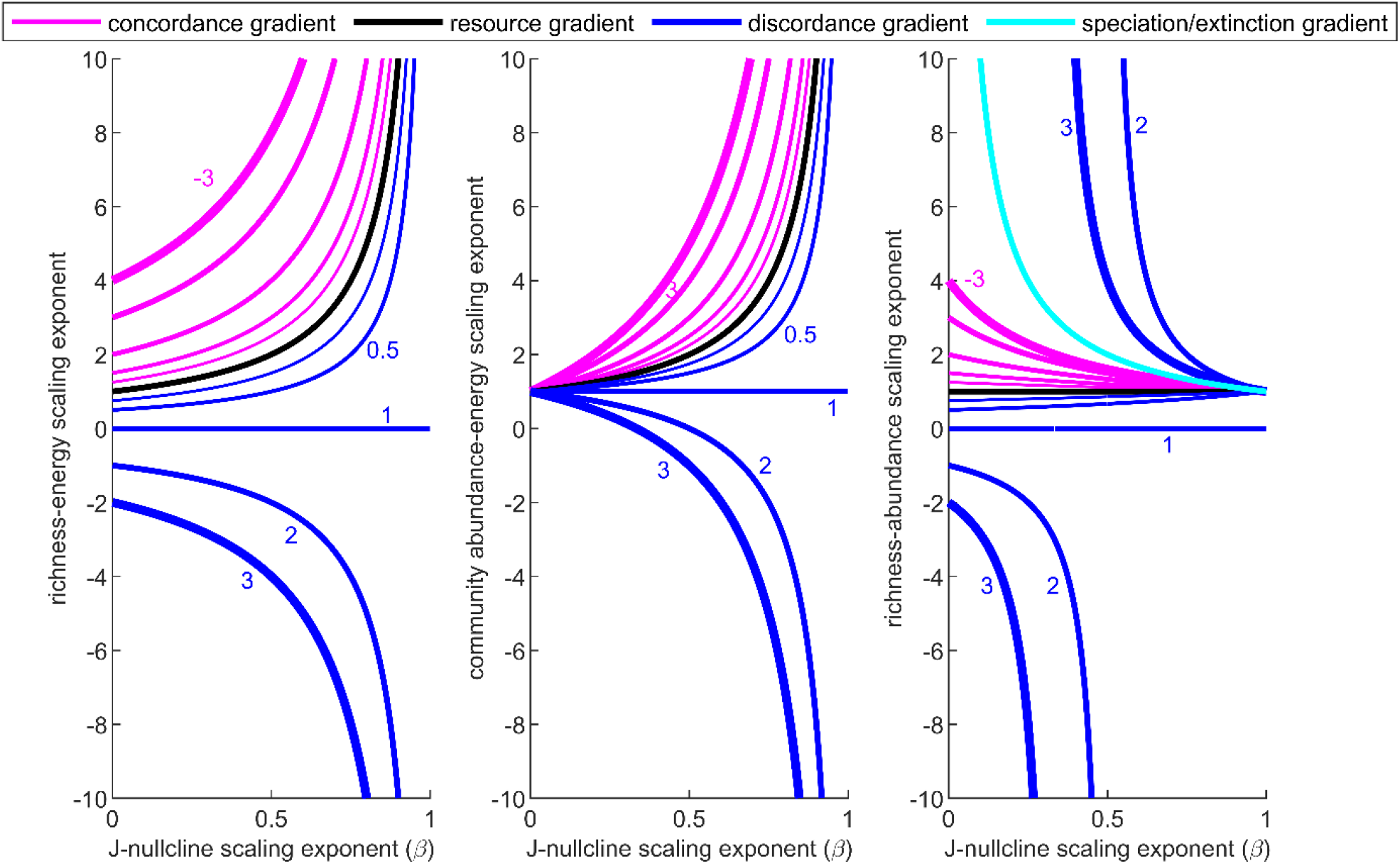
The predicted fffects of the J-nullcline on the scaling of the equilibria of species richness (*S*) and community abundance (*J*) along simple and complex environmental gradients. *β* is the scaling exponent of the J-nullcline, which quantifies the biodiversity effect on community abundance (BECA). The curves indicate the *S*-*J*-*E* scaling exponents for gradients in which speciation/extinction and resource availability covary in different ways. In a complex gradient, equilibrium mean population size 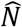 may decrease with resource availability due to higher speciation or lower extinction in high resource environments (a *concordance gradient*, pink curves,) or increase with resources (*a discordance gradient*; dark blues curves). In a concordance gradient, the scaling of *S* with *J* has a value between 1, the value expected for a pure gradient in energy availability, and 1/*β*, the scaling expected for a pure speciation/extinction gradient. Thicker curves indicate gradients having larger absolute values of *g*_*1*_/*r*_*1*_, where *r*_*1*_ quantifies the degree to which resource availability changes along the gradient and *g*_*1*_ quantifies the degree to which equilibrium mean abundance change along the gradient (Box 1). The value of *g*_*1*_/*r*_*1*_ is shown next to each curve.

When speciation/extinction and energy availability are changing as a power function of *G* (i.e., that 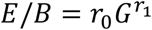 and 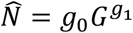), ETBD predicts:

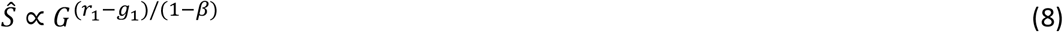

and

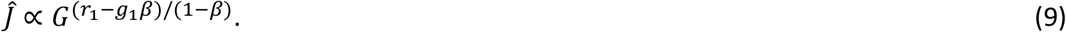

When speciation/extinction and energy availability are changing as an exponential function of *G* (i.e., 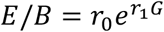 and 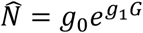), ETBD predicts:

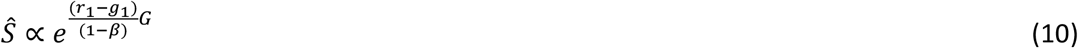

And

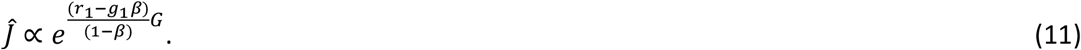

These equations demonstate that the biodiversity effect on community abundance (BECA), as quantified by *β*, shapes patterns of diversity and community abundance along environmental gradients such as temperature and latitude, pointing to a need to incorporate BECA into other theories and models of biodiversity, such as the metabolic theory of ecology and the neutral theory (Allen et al., 2002; Hubbell, 2001).

